# Disrupted MOS signaling alters meiotic cell cycle regulation and the egg transcriptome

**DOI:** 10.1101/2025.04.02.646881

**Authors:** Gisela Cairo, Olha Kholod, Olivia Palmer, Brittany A. Goods, Soni Lacefield

## Abstract

Mammalian female meiosis is tightly regulated to produce a developmentally competent egg. Oocytes enter meiosis in the fetal ovary and then arrest at prophase I until sexual maturation. Upon hormonal stimulation, a subset of oocytes resumes meiosis at which time, new transcription is halted. Oocytes then complete meiosis I, enter metaphase II, and arrest until fertilization, a process essential for egg competency and fertility. The MOS kinase is a key regulator of the metaphase II arrest, activating the MAPK signaling cascade. Loss of MOS in female mice disrupts the maintenance of the metaphase II arrest, leading some eggs to extrude two polar bodies and some to divide beyond anaphase II. To investigate the consequences of the *Mos* deletion, we performed live imaging and found that *mos^-/-^* eggs exhibit transient chromosome separation events in meiosis I, suggesting a role for MOS in coordinating the timing of meiotic divisions. Further analysis showed that new transcription is required for *mos^-/-^* eggs to undergo additional divisions but not for second polar body extrusion. Surprisingly, single-egg sequencing revealed extensive differences in gene expression between wildtype and *mos^-/-^*eggs, including those with only one polar body. Many of the differentially expressed genes were involved in cell cycle regulation, including *Aurka*, *Bub3*, and *Cdk7*. Other upregulated pathways included metabolism of RNA, transcription, and neddylation. Furthermore, the gene expression profile of *mos^-/-^* eggs was markedly different from that of wildtype eggs chemically activated to undergo embryo-like divisions. Our findings demonstrate that MOS plays a crucial role in meiotic cell cycle regulation and helps ensure that the egg maintains the proper transcriptome necessary for developmental competence.

## INTRODUCTION

In female mammals, meiosis is uniquely regulated and takes place within the specialized process of oogenesis to ensure the production of a developmentally competent egg. In the fetal ovary, primordial germ cells differentiate into oogonia, which eventually enter meiosis, undergo DNA replication, and arrest as oocytes at prophase I (Pan and Li, 2019). At sexual maturation, hormonal cues trigger a subset of oocytes to enter the first meiotic division, in which half the chromosomes are extruded into a polar body. The metaphase II spindle then reassembles, at which time the oocyte transitions into a developmentally competent egg. Meiosis II is only completed after fertilization, when the sperm triggers the segregation of sister chromatids, with half extruded into a second polar body. Understanding the molecular regulators of meiosis in oocytes is critical for preserving reproductive fidelity, as errors in meiosis can lead to infertility, early embryo loss, or the development of germ cell tumors.

Meiosis is tightly regulated to ensure the proper timing and coordination of cell cycle events. As oocytes resume the meiotic divisions, new transcription is halted, and protein production is dependent on translational control (Christou-Kent *et al*., 2020). Two major cell-cycle regulators, cyclin B and the MOS (Moloney Sarcoma Oncogene) kinase are translated as oocytes resume the meiotic divisions (Gebauer *et al*., 1994; Gebauer and Richter, 1997; Kotani *et al*., 2013). Cyclin B binds and activates cyclin-dependent kinase 1 (CDK1), the major cell cycle kinase that governs meiosis I and meiosis II chromosome segregation by phosphorylating specific substrates (Kim *et al*., 2023). As oocytes enter metaphase I, cyclin B levels rise, activating CDK1 for phosphorylation of substrates involved in spindle assembly and kinetochore-microtubule attachments. CDK1-cyclin B also phosphorylates and activates its own inhibitor, the Anaphase Promoting Complex/ Cyclosome (APC/C), a ubiquitin ligase that targets cyclin B for proteasomal degradation (Pines, 2011). Upon APC/C activation at anaphase I onset, cyclin B and other substrates are degraded and chromosomes segregate (Kim *et al*., 2023). CDK1 activity declines, allowing spindle disassembly. For oocytes to enter metaphase II, the APC/C is inhibited, cyclin B levels rise, and the metaphase II spindle assembles. The egg then arrests to await fertilization.

The inhibition of APC/C maintains metaphase II arrest in mice through two pathways. First, a direct inhibitor, EMI2, binds APC/C and brings the phosphatase PP2A to dephosphorylate APC/C components (Hörmanseder *et al*., 2013). Second, the MOS kinase activates a MAPK (mitogen-activated protein kinase) signaling cascade to maintain APC/C inhibition. Loss of EMI2 prevents mouse oocytes from entering meiosis II due to unrestrained APC/C activation that results in low cyclin B levels (Madgwick *et al*., 2006; Shoji *et al*., 2006; Pasternak *et al*., 2016). In contrast, *mos^-/-^* eggs enter metaphase II, but most cannot maintain the arrest (Colledge *et al*., 1994; Hashimoto *et al*., 1994). Some *mos^-/-^* eggs escape metaphase II, progress through anaphase II and then arrest, while others continue dividing beyond anaphase II. The additional divisions give rise to an embryo-like structure, which cannot develop into a viable organism. This embryo-like mass of cells was originally termed as a parthenogenone, which refers to a type of parthenote made up of haploid cells, originally derived from the activation of an unfertilized haploid egg (Colledge *et al*., 1994). We will call them parthenotes because we do not know the ploidy of all cells within the mass. Furthermore, approximately one-third of *mos^-/-^*mice develop ovarian teratomas, suggesting that a subset of the activated eggs that form parthenotes can eventually undergo further uncontrolled divisions to become malignant tumors (Colledge *et al*., 1994; Hashimoto *et al*., 1994; Hirao and Eppig, 1997).

Although MOS expression is restricted to oocytes, specifically during the meiotic divisions, the loss of MOS has significant consequences for embryo development. Variants of *MOS* have been identified through whole exome sequencing from patients in fertility clinics (Zhang *et al*., 2021, 2022; Jiao *et al*., 2022). In vitro fertilization (IVF) of the eggs resulted in uncontrolled divisions, causing early embryonic arrest and fragmentation. These results highlight that infertility in these patients is likely due to early embryo loss and cannot be treated through IVF alone. Additionally, *mos^-/-^* female mice also have significantly reduced fertility, suggesting that the eggs are not able to produce a viable embryo upon fertilization (Colledge *et al*., 1994; Hashimoto *et al*., 1994). These results suggest that even the eggs that appear normal, with one polar body, are likely not competent for normal embryo development.

MOS/MAPK signaling is known to have several roles in meiosis, including in the regulation of spindle assembly, cortical actin thickening, and mRNA translation and degradation (Choi *et al*., 1996; Verlhac *et al*., 1996, 2000; Lefebvre *et al*., 2002; Terret *et al*., 2003; Deng *et al*., 2005; Chaigne *et al*., 2015; Sha *et al*., 2017; Kalous *et al*., 2018; Cao *et al*., 2020). However, whether MOS has additional functions in cell cycle regulation remains unclear, particularly in relation to how eggs that maintain one polar body differ from those that undergo additional divisions. In this study, we aimed to further define the role of MOS in meiotic regulation and investigate how loss of MOS leads to developmentally incompetent eggs and the potential for abnormal, additional cell divisions. Using the mouse model system, we found that MOS temporally regulates meiosis I chromosome segregation. Furthermore, we show that new transcription is required for *mos^-/-^*eggs to undergo additional cell divisions. Interestingly, however, using single oocyte and single parthenote sequencing, we found that not only the parthenotes, but also *mos^-/-^* eggs with one and two polar bodies also underwent new transcription. The differential gene expression analysis showed that *mos^-/-^* eggs had a markedly distinct pattern of expression from both wildtype eggs arrested at metaphase II and from wildtype eggs that were chemically activated to undergo parthenogenesis. Our results showed an increased expression in maternal mRNAs, embryonic genes and many other genes, including those involved in cell-cycle regulation. These results suggest that *mos^-/-^* eggs undertake an abnormal gene expression pattern that likely contributes to the formation of eggs that are not competent for normal embryo development after fertilization, leading to infertility.

## RESULTS

### A subset of *mos^-/-^* eggs undergo transient chromosome separation events prior to the first polar body extrusion

While the MOS/MAPK signaling pathway has a well-established function in maintaining metaphase II arrest, we first wanted to assess whether MOS had additional roles in meiotic regulation (Colledge *et al*., 1994; Hashimoto *et al*., 1994). Based on previous studies that used immunofluorescence images to study *mos^-/-^*mutants, we hypothesized that time-lapse imaging could uncover additional abnormalities in meiosis. To this end, we collected prophase I arrested oocytes from wildtype and *mos^-/-^*mutants, released them from the arrest, and initiated time-lapse imaging. The cell stains SYP555-tubulin and SiR-DNA were added to monitor spindles and DNA, respectively (Fig. 1A-C). As expected, wildtype oocytes underwent a first meiotic division in which half the chromosomes were extruded into a polar body and then they arrested with a metaphase II spindle (Fig. 1A).

**Figure 1.**
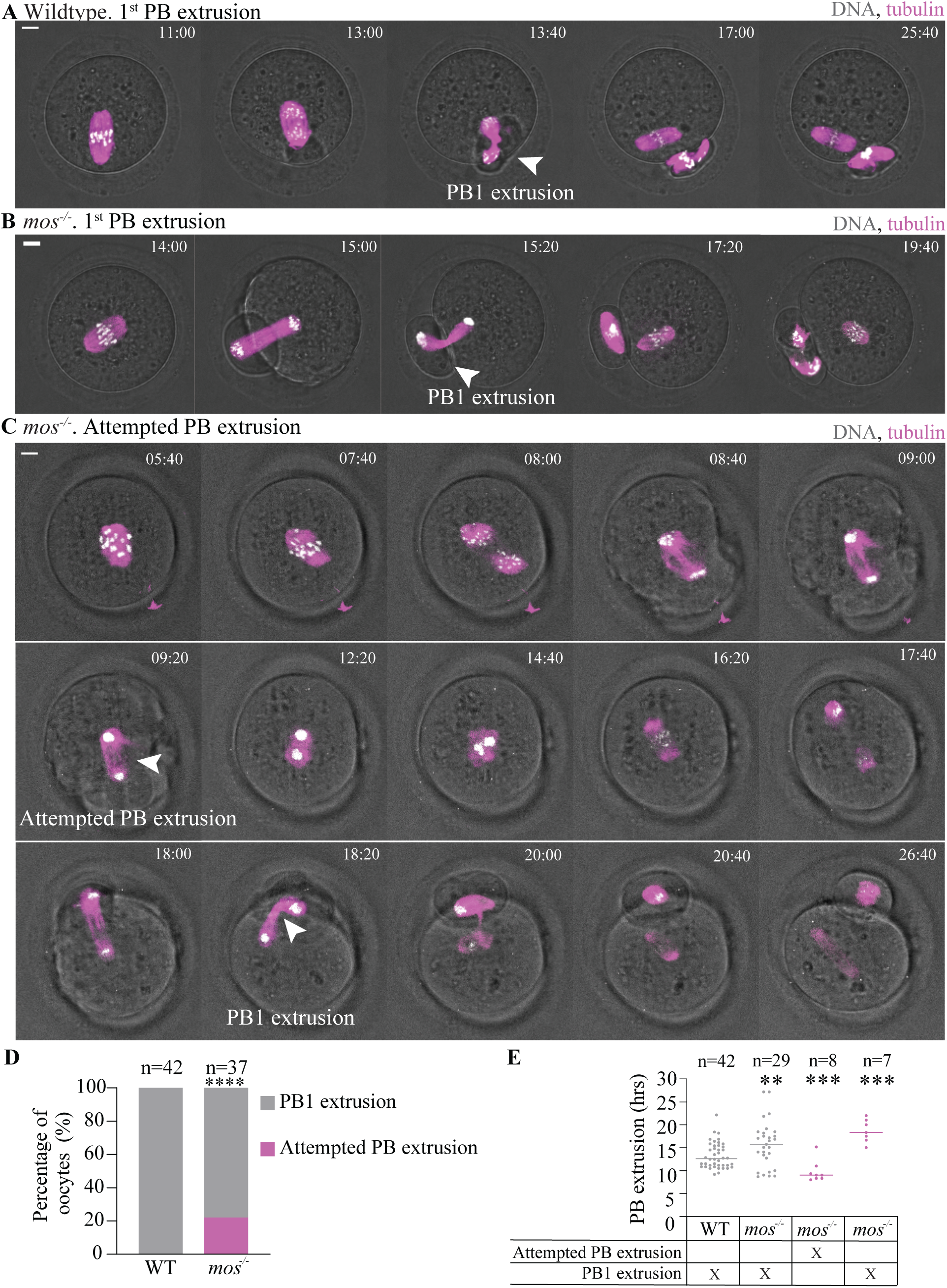
**Some mo*s^-/-^* eggs undergo transient chromosome segregation events**. **(A-C)** Representative time-lapses of wildtype (A) and *mos^-/-^* (B-C) oocytes undergoing the first polar body extrusion (A-B) or attempted polar body extrusion before undergoing a complete polar body extrusion (C). SPY555tubulin and SirDNA was used to visualize tubulin (magenta) and DNA (white), respectively. Time 0h corresponds to germinal vesicle breakdown. White arrows indicate first PB extrusion or attempted PB extrusion. Scale bars, 10µm. **(D)** Percent of wildtype and *mos^-/-^* oocytes that undergo attempted polar body extrusions or complete polar body extrusions on the first round of chromosome segregation (n indicates the number of oocytes per genotype). 3 or more independent experiments were conducted per genotype. Asterisks indicate a statistically significant difference compared with wildtype (****, p < 0.0001, two-tailed Fisher’s exact test). **(E)** Graph of the time to first polar body extrusion or attempted extrusion in wildtype and *mos^-/-^* oocytes. Time 0h corresponds to germinal vesicle breakdown. Grey data points correspond to wildtype or *mos^-/-^*eggs that extrude the first PB; pink data points correspond to *mos^-/-^* eggs that attempt polar extrusion before a completed polar body extrusion (****, p < 0.05, Mann-Whitney test). WT, wildtype; PB, polar body; PB1, first polar body.

Surprisingly, ∼22% of *mos^-/-^* oocytes attempted chromosome segregation and polar body extrusion but then retracted the polar body and brought the spindle and chromosomes back together (Fig. 1C-D). This unsuccessful attempt is followed by another chromosome segregation event, leading to a permanent polar body extrusion. The first attempt at chromosome segregation in *mos^-/-^* mutants occurs on average 10 hours after prophase I release, which is on average 3 hours earlier than the first chromosome segregation in wildtype oocytes (Fig. 1E). In *mos^-/-^*mutants, the timing of the onset of permanent chromosome segregation is variable but occurs on average 4 hours after wildtype oocytes. These chromosome segregation attempts could not be revealed from single-timepoint images, but time-lapse imaging gives a more complete understanding of the abnormalities. Overall, our data demonstrate that *mos^-/-^* mutants have previously undescribed defects in meiotic regulation, with additional chromosome segregation and polar body extrusion attempts and a delayed completed polar body extrusion.

### *mos^-/-^* mutant eggs undergo a variable number of cell divisions after spontaneous activation

Previous studies showed that a proportion of *mos^-/-^*eggs spontaneously activate and undergo additional cell divisions after meiosis II (Colledge *et al*., 1994; Hashimoto *et al*., 1994). To further understand the variability in the cellular events that occur in individual *mos^-/-^* eggs, we used time-lapse imaging to monitor the additional cell divisions. Because *mos^-/-^* mutants undergo spontaneous parthenogenetic activation, we compared *mos^-/-^* eggs to both untreated wildtype eggs and to wildtype eggs that were chemically activated to undergo parthenogenesis with the addition of strontium chloride (SrCl_2_) (Jellerette *et al*., 2000). Chemical activation mimics fertilization, triggering processes like maternal mRNA processing, increased protein synthesis, mitochondrial activity for ATP production, and membrane dynamics such as cortical granule exocytosis (Jellerette *et al*., 2000; Horner and Wolfner, 2008; Krauchunas and Wolfner, 2013; Bellido-Quispe *et al*., 2024). We isolated *mos^-/-^*and wildtype prophase I arrested oocytes, released them from the arrest in culture and then initiated the imaging approximately 13-15 hours after the release, a timepoint in which the oocytes had undergone the first polar body extrusion (Fig. 2A-D). The chemically activated wildtype eggs were treated with SrCl_2_ for 2-3 hours prior to initiating the imaging (Fig. 2A).

**Figure 2.**
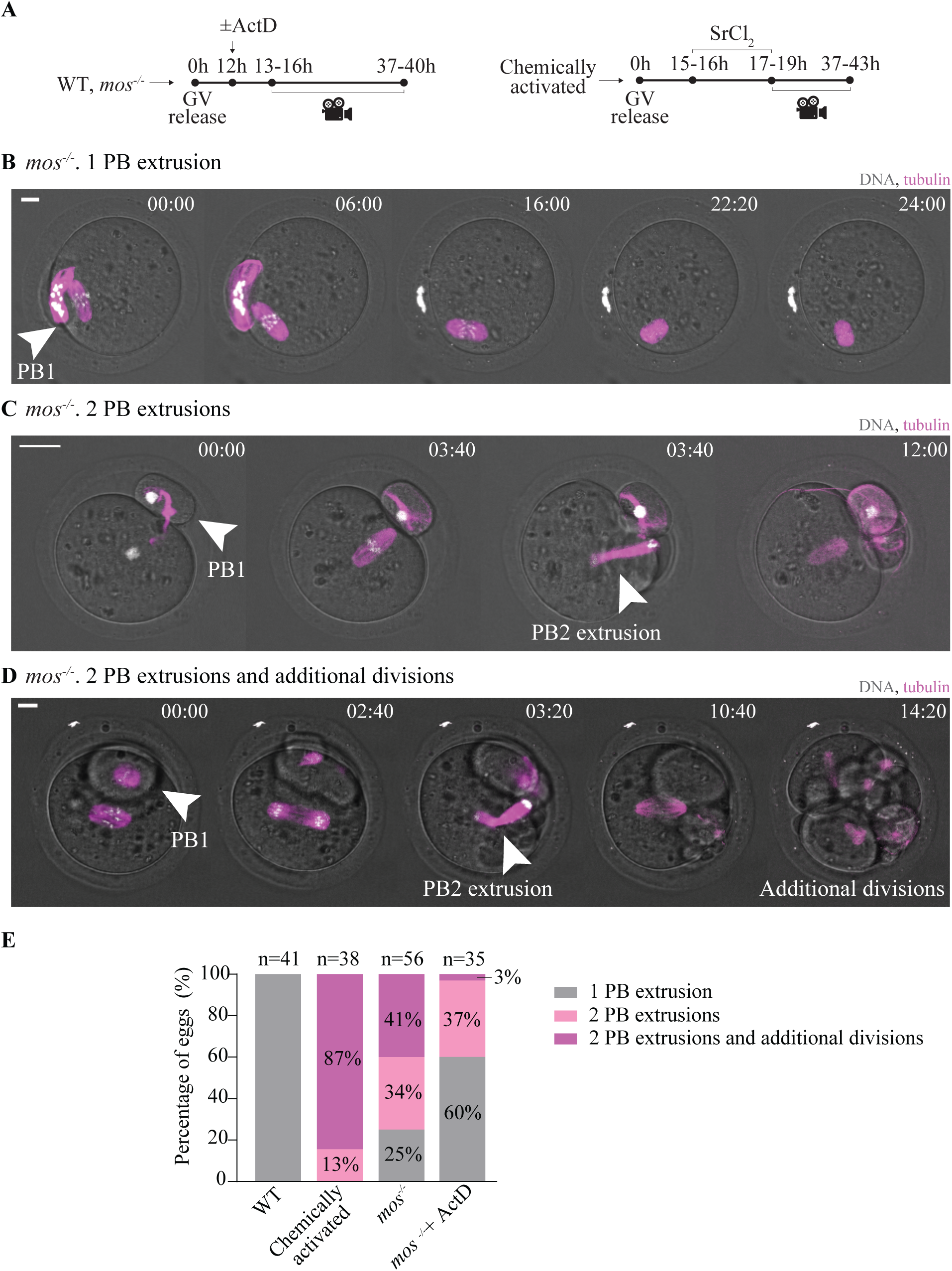
***mos^-/-^*eggs show variable number of polar body extrusions and cell divisions after spontaneous activation**. **(A)** Schematic of experimental setup. Wildtype and *mos^-/-^* oocytes were imaged starting 13-15h after meiotic resumption (GV release) for 24 hours. If applicable, actinomycin was added 12h after GV release. Chemically activated wildtype eggs were incubated with SrCl_2_ for 2-3 hours and subsequently imaged for 24 hours, starting 15-16 hours after GV release. **(B-D)** Representative time-lapse images of *mos^-/-^*eggs undergoing 1 PB extrusion (B), 2 PB extrusions (C), and 2 PB extrusions and additional divisions (D). SPY555tubulin and SirDNA were added to visualize tubulin (magenta) and DNA (white), respectively. Time 0 represents the start of live imaging, following extrusion of the first PB (B-D). White arrows indicate the time of 1^st^ or 2^nd^ PB extrusion. Scale bars, 10µm. **(E)** Percent of eggs that extruded 1 PB, 2 PBs, or 2 PBs and additional divisions. n indicates the number of eggs imaged per condition. 3 or more independent experiments were conducted per genotype/ condition. GV, germinal vesicle; WT, wildtype; PB, polar body; ActD, actinomycin D.

As expected, wildtype eggs arrested at metaphase II after extruding a single polar body (Fig. 2E). All chemically activated eggs underwent parthenogenetic activation, extruding a second polar body, and 87% proceeded to undergo at least one additional cell division (Fig. 2E). Strikingly, we observed variability in *mos^-/-^* eggs, with 25% extruding one polar body, 34% extruding 2 polar bodies, and 41% extruding 2 polar bodies and then undergoing at least one additional division (Fig. 2B-E). In addition to the live imaging, we observed the same phenotypes with immunostaining of wildtype and *mos^-/-^* eggs 24 hours after prophase I release, imaging tubulin or actin along with DNA (Figure S1A-D). These results demonstrated that the phenotypes observed were not due to phototoxicity or the use of the dyes. Overall, our results align with previous studies demonstrating that *mos^-/-^* eggs exhibit variable phenotypes (Colledge *et al*., 1994; Hashimoto *et al*., 1994).

The advantage of using live imaging is that we can follow the dynamic processes occurring in each individual egg. A previous report using immunofluorescence images reported that *mos^-/-^* eggs arrest for 2-4 hours prior to resuming meiosis (Verlhac *et al*., 1996). We further analyzed the timing of metaphase II exit to further understand the variability in individual eggs in relation to the phenotypic outcome (Fig. 3A-D). Wildtype oocytes stayed arrested at the metaphase II arrest for the duration of the imaging (longer than 15 hours) (Fig. 3A, D). In contrast, chemically activated wildtype eggs released from the metaphase II arrest within 0-4 hours after SrCl_2_ induction (Fig. 3D). The *mos^-/-^* mutants exhibited variability in the time of the arrest (Fig. 3B-D). Of those that extruded one polar body, 36% disassemble their spindles and enter an interphase state, without undergoing anaphase II (Fig. 3B, D). The remaining 64% arrested at metaphase II for the duration of the imaging (Fig. C-D). The *mos^-/-^* eggs that undergo 2 polar body extrusions only stayed in metaphase II 0-4hrs before undergoing anaphase II (Fig. 3D). Similarly, those eggs that underwent additional divisions after metaphase II also only remained in metaphase II for 0-4hrs before undergoing anaphase II. These results suggest that the eggs that parthenogenetically activate will exit metaphase II quickly, supporting prior reports (Verlhac *et al*., 1996). Our data further suggest that many of the *mos^-/-^* eggs that undergo one polar body extrusion exit the cell cycle without completing anaphase II. This finding suggests that many of the *mos^-/-^* eggs with one polar body are abnormal because they cannot maintain the metaphase II arrested state, which is important for maintaining developmental competence.

**Figure 3.**
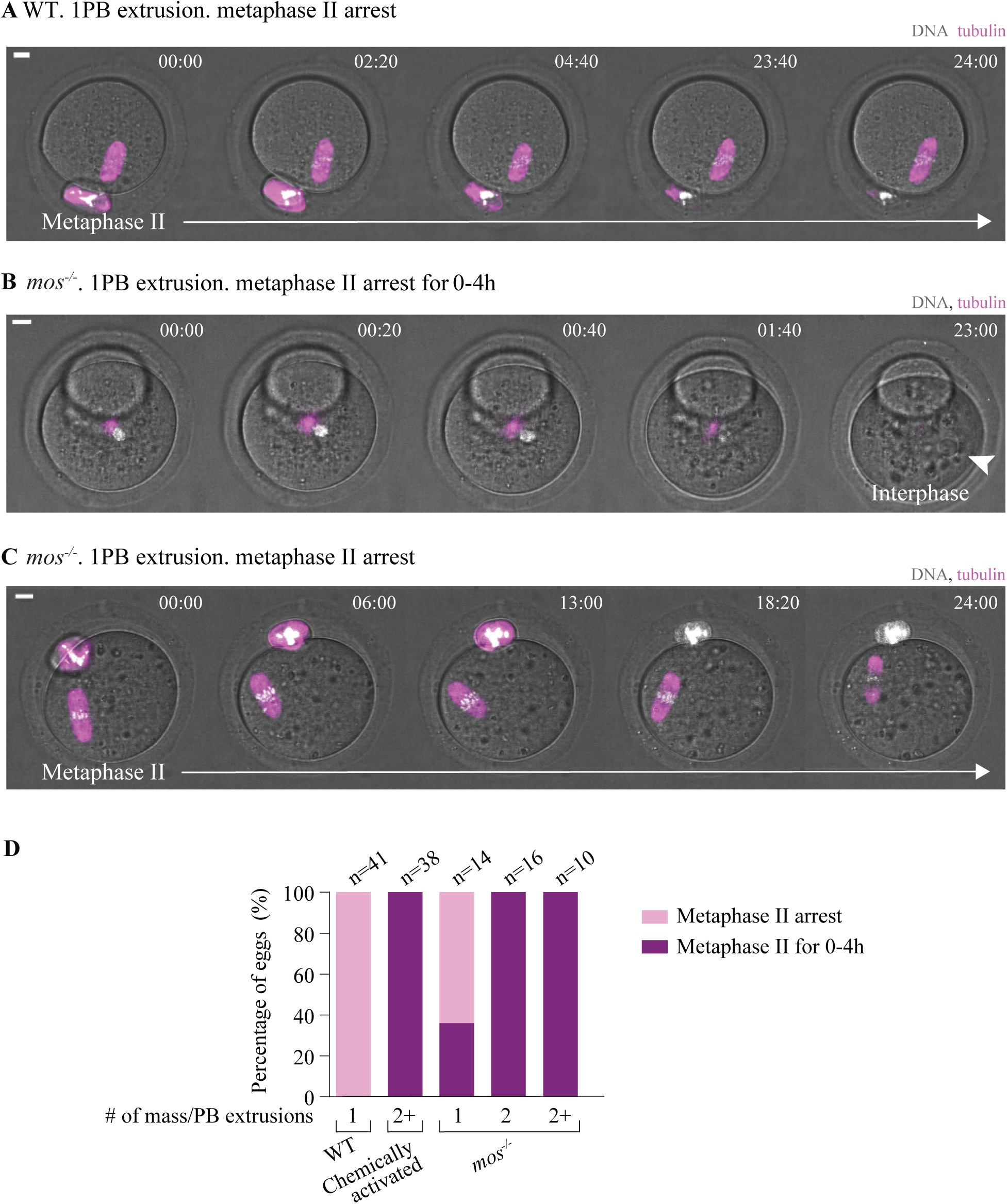
**Metaphase II duration varies in *mos^-/-^* eggs that extrude 1 PB**. **(A-B)** Representative time-lapse images of *mos^-/-^* eggs at metaphase II arrest for the indicated time. Wildtype and *mos^-/-^* eggs were imaged starting 13-15h after GV release for 24 hours. Chemically activated wildtype eggs were incubated in SrCl_2_ for 2-3 hours and subsequently imaged for 24 hours, starting 15-16 hours after GV release. SPY555tubulin and SirDNA was added to visualize tubulin (magenta) and DNA (white), respectively. Time 0 represents the start of live imaging. Scale bars, 10µm. **(C)** Percent of wildtype, chemically activated, and *mos^-/-^* eggs that undergo metaphase II arrest within these time intervals: 0 to 4 hours, or for the duration of the movie, which were classified as metaphase II arrest. The eggs are further classified based on the number of PBs extruded and additional divisions. 3 or more independent experiments were conducted per genotype/condition. ActD, actinomycin; WT, wildtype; PB, polar body; 2+, 2 PB extrusions and additional divisions.

### New transcription is required for *mos^-/-^* eggs to undergo additional divisions

In wildtype oocytes, transcription is halted throughout meiosis, including during metaphase II arrest (Christou-Kent *et al*., 2020). New transcription only occurs after fertilization, in late one-cell and early two cell embryos (Aoki, 2022). Given that new transcription is needed for the early embryonic mitotic cell cycles, we hypothesized that the *mos^-/-^* eggs that undergo additional divisions may inappropriately activate embryonic gene expression without fertilization. To determine if transcription is required for the additional divisions, we added the transcription inhibitor actinomycin D to *mos^-/-^* oocytes in culture after they extruded the first polar body (Fig 2A). The live imaging revealed that 60% of these eggs with one polar body remained the same, 37% extruded a second polar body, and only 3% underwent additional divisions (Fig. 2E). If we compare these results to *mos^-/-^*eggs without actinomycin D treatment, we find a similar proportion that extruded two polar bodies. In contrast, the *mos^-/-^* eggs with actinomycin D treatment have a greater proportion that only extrude one polar body and much fewer with additional divisions when compared to *mos^-/-^*eggs without treatment. These results suggest that the inhibition of transcription prevented a proportion of *mos^-/-^*eggs from extruding a second polar body and undergoing additional divisions. Furthermore, our data demonstrate that new transcription is required for *mos^-/-^* eggs to undergo multiple divisions. Although the results support our hypothesis, we were surprised by this finding because new transcription is halted throughout meiosis, suggesting that the eggs must enter a different cell cycle state after parthenogenetic activation.

### Wildtype, chemically activated, and *mos^-/-^* eggs have distinct gene expression patterns

Our finding that new transcription is needed for *mos^-/-^* eggs to undergo multiple divisions led us to ask which specific transcripts are dysregulated and if early embryonic genes are inappropriately expressed in the eggs that underwent multiple divisions. To address this question, we decided to use single egg and parthenote RNA-sequencing (RNA-seq). We first used our live imaging to identify a timepoint in which we could classify the phenotypes (Fig. 4A-C). At 38-40h in culture, wildtype eggs had one polar body, while the chemically activated wildtype eggs underwent at least one division after anaphase II; *mos^-/-^* eggs had the expected varied phenotypes that we further classified, some with one polar body (M1), two polar bodies (M2), and some that underwent multiple divisions and became parthenotes (M3) (Fig. 4A-C).

**Figure 4.**
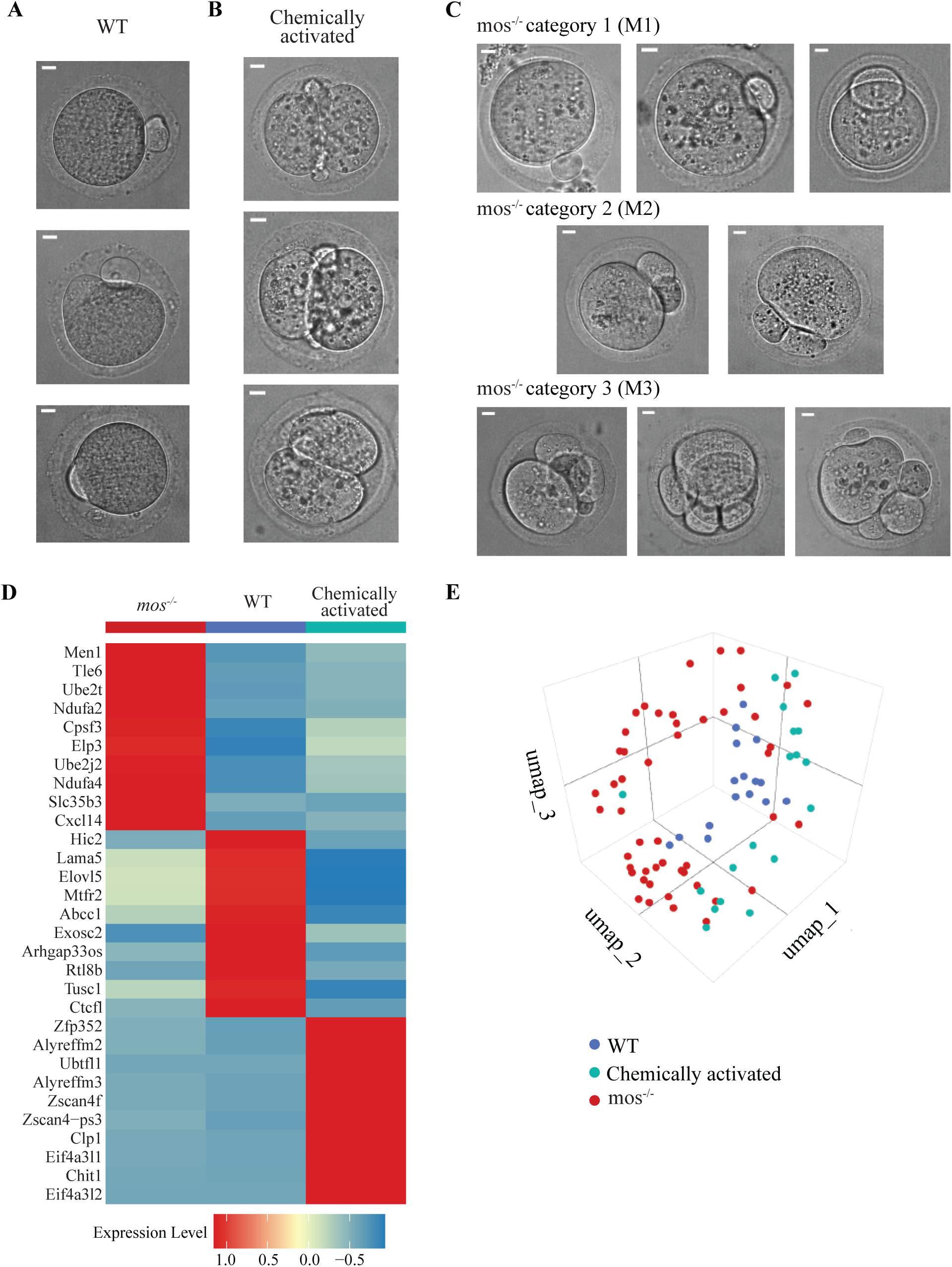
**Wildtype, chemically activated, and *mos^-/-^* eggs differentially express genes**. **(A-C)** Representative widefield images of wildtype (A), chemically activated (B), and *mos^-/-^* eggs (C) from time-lapse imaging taken 38-40h after GV release. *mos^-/-^* eggs are classified into the categories M1 (1PB), M2 (2PBs), and M3 (2PBs and additional divisions). Scale bars, 10µm. **(D)** Heatmap displaying the top 10 differentially expressed genes, comparing wildtype, chemically activated wildtype, and *mos^-/-^* eggs. Each row corresponds to a specific gene, while each column represents the average expression level for that gene. **(E)** Three-dimensional UMAP projection of scRNA-seq data (n = 107) colored by class: WT (blue), chemically activated (green), and *mos^-/-^* eggs (red). WT, wildtype; PB, polar body.

We then performed an experiment in which we isolated single oocytes from wildtype and *mos^-/-^* mutants, released them from the prophase I arrest, and matured them for 38-40h in culture, took an image under the stereoscope, and isolated single cells for RNA-sequencing (RNA-seq). We used well-established Smart-Seq2 protocols to generate RNA-seq libraries and sequenced 108 single eggs and parthenotes (Trombetta *et al*., 2014). Overall, we observed high quality sequencing data for each egg and parthenote, with a high number of unique reads detected per egg/parthenote (10 million), and a low mapping rate to rRNA (<20%), and >70% alignment rate to the genome (Fig. S2A-C).

Comparisons of differentially expressed genes showed an up-regulation of distinct transcripts when comparing wildtype, chemically activated, and *mos^-/-^* eggs and parthenotes, with significant highly increased transcripts in each category. The top 10 differentially expressed genes are shown for each category (Fig. 4D). Upon further analysis, a comparison of gene expression patterns between cells is displayed by Uniform Manifold Approximation and Projection (UMAP), with each sequenced egg represented (Satija *et al*., 2015; McInnes *et al*., 2020). This analysis highlights distinctions between individual wildtype, chemically activated, and *mos^-/-^* eggs (Fig. 4E). Overall, these results suggest that *mos^-/-^* eggs have unique gene expression patterns in comparison to both wildtype and chemically activated eggs.

A pairwise comparison between wildtype and *mos^-/-^* eggs and parthenotes identified 2793 of differentially expressed genes (DEGs) (p value < 0.05; log2foldchange >1 for upregulated or <-1 for downregulated) with the majority increased in expression in *mos^-/-^* eggs (2,759 upregulated and 34 downregulated; Fig. 5A, Supplementary Table 1). Metascape pathway analysis of the upregulated DEGs revealed molecular functions related to metabolism of RNA, transcription, cell-cycle regulation, neddylation, and cell signaling (Zhou *et al*., 2019)(Fig. 5B). The cell cycle pathways included G2/M transition, cell cycle checkpoints, and APC/C mediated degradation of cell cycle proteins. These results suggest that the *mos^-/-^* eggs and parthenotes have a unique cell cycle profile, which likely allowed them to enter abnormal divisions.

**Figure 5.**
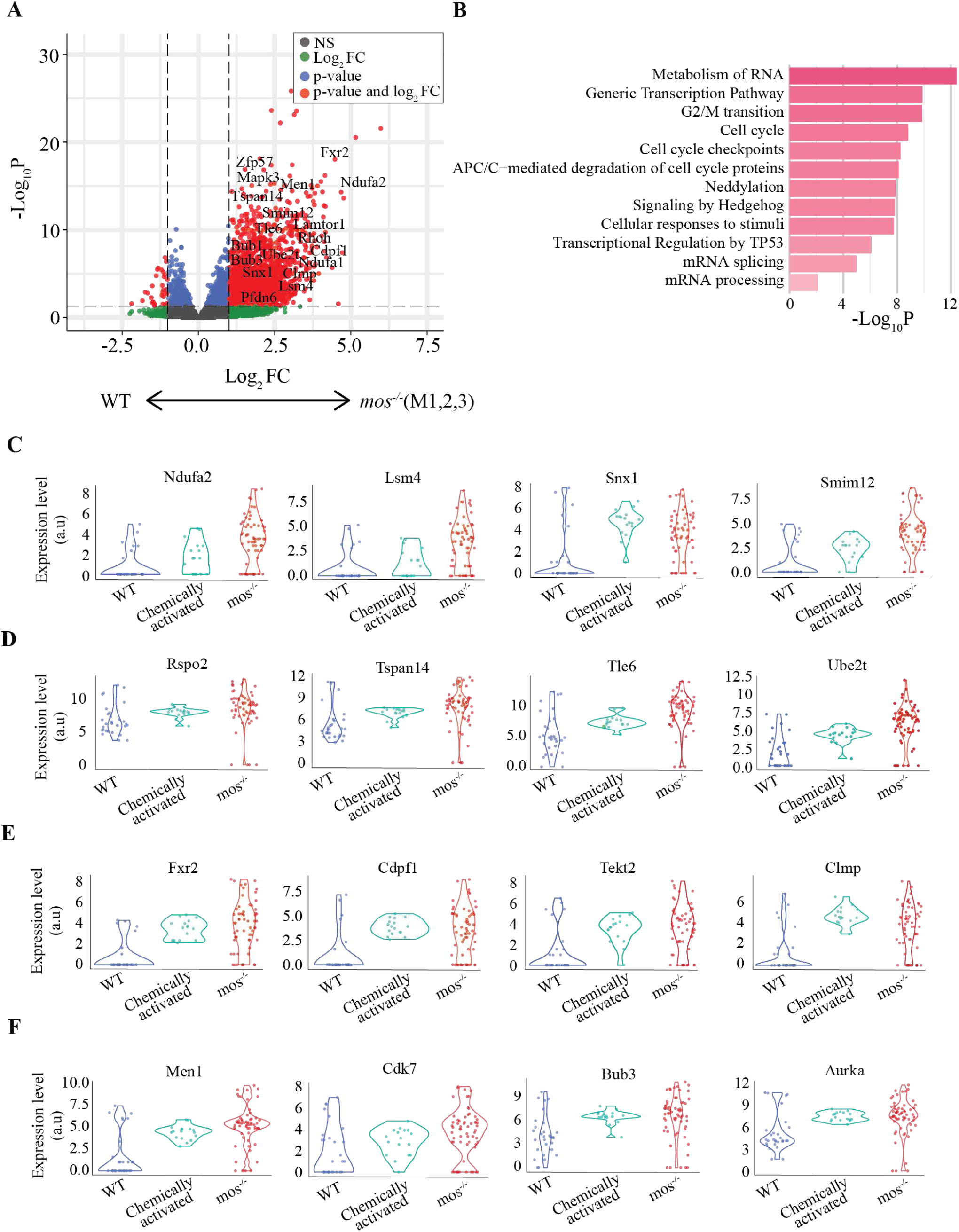
***mos^-/-^*eggs express genes associated with RNA metabolism, transcription, and cell cycle regulation**. **(A)** Volcano plot showing differentially expressed genes between *mos^-/-^* and wildtype eggs. Genes significantly upregulated in *mos^-/-^* eggs appear on the right, downregulated on the left, and non-significant changes are centered. All *mos^-/-^*eggs (M1, M2, M3) are represented in this dataset. **(B)** Bar plot depicting pathway enrichment analysis for genes upregulated in *mos^-/-^*eggs compared to wildtype eggs. Each bar represents a specific biological pathway, with the length indicating the degree of enrichment significance. **(C-F)** Violin plots representing upregulated genes in *mos^-/-^* eggs when compared to wildtype eggs. Each point is an individual egg/ parthenote. The genes were classified into categories: those normally induced during early embryogenesis (C); maternal RNAs typically degraded in the egg (D); genes whose transcript levels remain consistent throughout metaphase II and early embryogenesis (E); and interesting genes involved in cell cycle regulation. WT, wildtype.

Our original hypothesis stated that early embryonic genes were inappropriately expressed in the *mos^-/-^* eggs and parthenotes. Our analysis confirmed this, revealing that there was indeed an increased expression of genes normally expressed in early embryogenesis in *mos^-/-^* eggs and parthenotes compared to wildtype, including *Ndufa2, Lsm4, Snx1*, and *Smim12* (Fig. 5C). However, there was also an increase in other genes as well. For example, the *mos^-/-^* eggs and parthenotes showed an increase in maternal mRNAs that are typically degraded in the early embryo, including *Rspo2, Tspan14, Tle6*, and *Ube2t* (Fig. 5D). There were also genes whose expression normally stays consistent throughout metaphase II and early embryogenesis that were upregulated in *mos^-/-^*eggs and parthenotes, including *Fxr2, Cdpf1, Tekt2*, and *Clmp* (Fig. 5E). Additionally, interesting genes involved in cell-cycle regulation also had increased expression in *mos^-/-^* eggs and parthenotes, including the tumor suppressor gene Men1, Cdk7, Bub3, and Aurka (Fig. 5F). Together, these results suggest that not only are the early embryonic genes upregulated in *mos^-/-^*eggs and parthenotes in comparison to wildtype, but many other genes are also upregulated.

We also compared *mos^-/-^* and chemically activated wildtype eggs and parthenotes, hypothesizing that they would have a similar gene expression pattern. In both cases, the eggs undergo parthenogenetic activation and additional divisions. However, when we first compared the expression patterns of the wildtype eggs to the chemically activated wildtype parthenotes, we noticed a completely different pattern of DEGs as compared to the *mos^-/-^* eggs and parthenotes versus wildtype eggs (Fig. 6A, 5A). In comparison, the chemically activated wildtype parthenotes have many both increased (2950) and decreased (858) transcripts when compared to wildtype, whereas the *mos^-/-^* mutants have mostly increased transcripts when compared to wildtype. The pathways of the increased transcripts in chemically activated wildtype parthenotes in comparison to wildtype eggs include RNA metabolism, membrane trafficking, translation, ribonucleoprotein complex biogenesis, and mRNA metabolic processes (Fig. 6B). A direct comparison of the transcripts in *mos^-/-^* versus chemically activated wildtype eggs/ parthenotes showed many DEGs, including 1011 up-regulated and 1021 down-regulated transcripts (Fig. 6C). The pathways that were upregulated in *mos^-/-^* eggs/parthenotes include processing pre-mRNA, metabolism of RNA, cell cycle, and chromatin organization (Fig. 6D). These results demonstrate that the *mos^-/-^* eggs/parthenotes have a distinct gene expression pattern from both wildtype eggs and chemically activated wildtype parthenotes.

**Figure 6.**
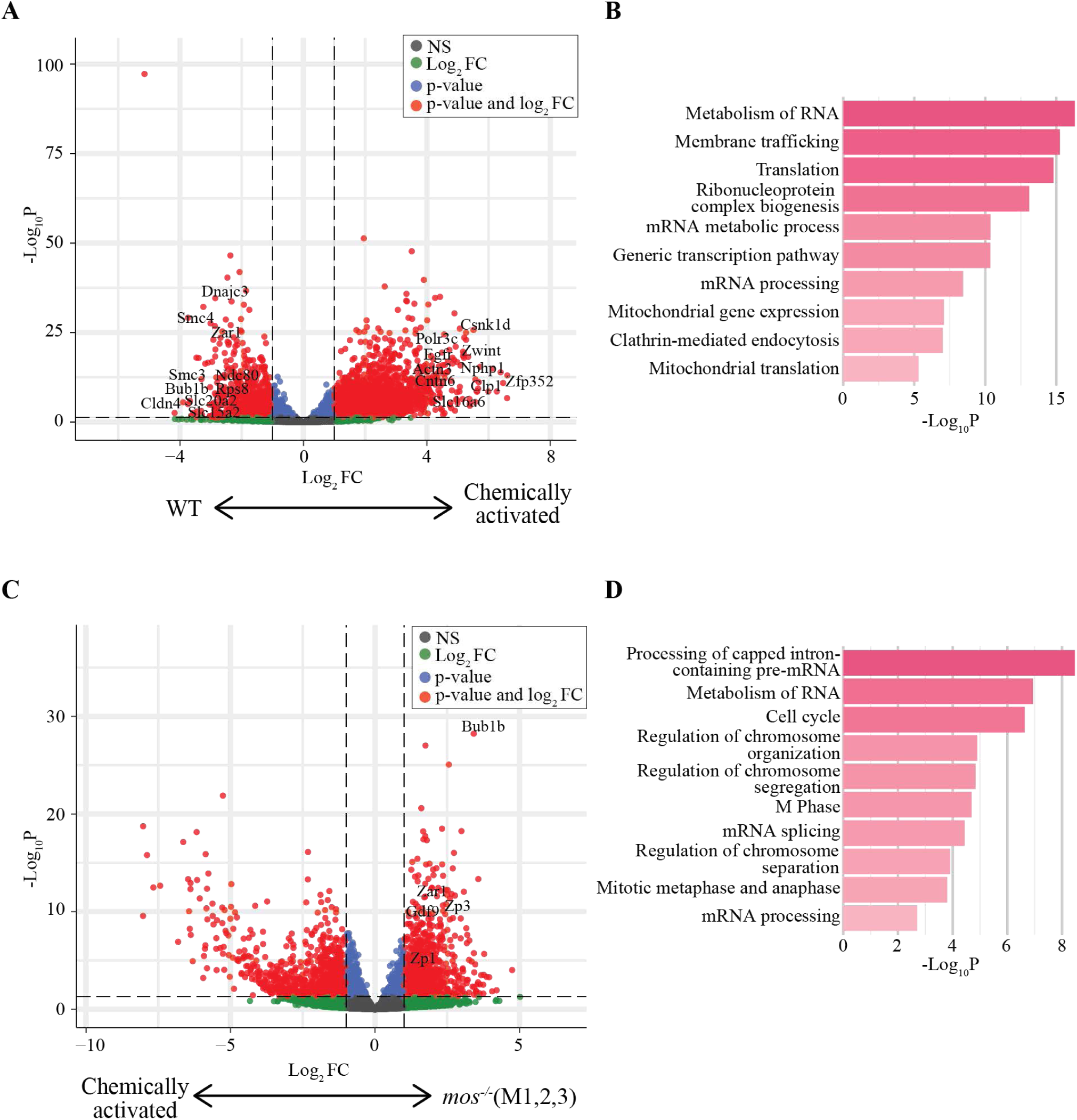
Wildtype, chemically activated, and *mos^-/-^* eggs have differentially expressed genes. **(A)** Volcano plot showing differentially expressed genes between untreated wildtype eggs (WT) and chemically activated wildtype eggs/parthenotes. Genes significantly upregulated in chemically activated eggs/parthenotes appear on the right, those downregulated on the left, and non-significant changes are centered. **(B)** Bar plot depicting pathway enrichment analysis for genes upregulated in chemically activated wildtype eggs compared to untreated wildtype eggs. Each bar represents a specific biological pathway, with the height indicating the degree of enrichment significance. **(C)** Volcano plot showing differentially expressed genes between chemically activated wildtype and *mos^-/-^* eggs/parthenotes (M1, M2, M3). Genes significantly upregulated in chemically activated eggs appear on the right, those downregulated on the left, and non-significant changes are centered. **(D)** Bar plot depicting pathway enrichment analysis for genes upregulated in *mos^-/-^*eggs/parthenotes compared to chemically activated eggs/parthenotes. Each bar represents a specific biological pathway, with the height indicating the degree of enrichment significance. WT, wildtype.

### *mos^-/-^* eggs with one polar body have increased gene expression when compared to wildtype

An advantage of sequencing single eggs and parthenotes is that we could link the DEGs with the phenotypic outcome of whether the *mos^-/-^* eggs had one polar body (M1), two polar bodies (M2), or two polar bodies and additional divisions (M3). Because our previous results showed that the inhibition of transcription strongly reduced the population of *mos^-/-^*eggs that underwent additional divisions beyond anaphase II (Fig. 2E), we had originally hypothesized that only the M3 population would have increased gene expression. However, the UMAP comparison of each egg or parthenote showed that M1, M2, and M3 categories did not fit into individual regions that were distinct from one another (Fig. 7A). These results suggest that the eggs and parthenotes may have a more similar gene expression pattern than we had hypothesized.

**Figure 7.**
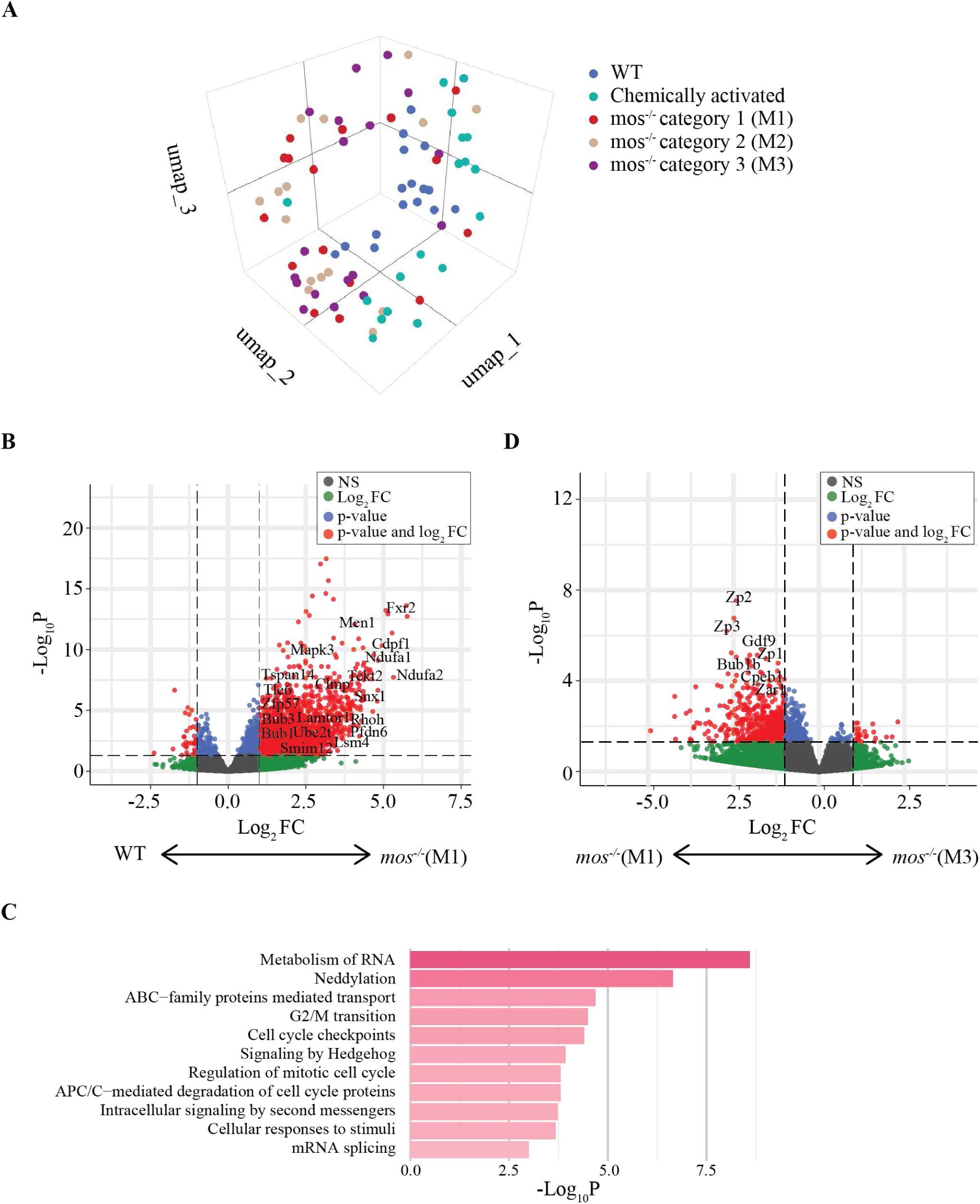
The *mos^-/-^*eggs with one polar body have many differentially expressed genes when compared to wildtype eggs. **(A)** Three-dimensional UMAP projection of scRNA-seq data (n = 108) colored by type: wildtype (blue), chemically treated (green), *mos^-/-^* eggs M1 (red), M2 (beige), or M3 (purple). M1 and M2 correspond to *mos^-/-^*eggs that extrude 1 or 2 PBs, respectively; M3 corresponds to *mos^-/-^*eggs that extrude 2 PBs and undergo additional divisions. **(B)** Volcano plot showing differentially expressed genes between *mos^-/-^* eggs that extrude 1 PB (M1) and wildtype eggs. Genes significantly upregulated in M1 eggs appear on the right, those downregulated on the left, and non-significant changes are centered. **(C)** Bar plot depicting pathway enrichment analysis for genes upregulated in *mos^-/-^*eggs that extrude 1 PB compared to wildtype eggs. Each bar represents a specific biological pathway, with the length indicating the degree of enrichment significance. **(D)** Volcano plot showing differentially expressed genes between *mos^-/-^*eggs that extrude 2 PBs and undergo additional divisions (M3) and *mos^-/-^* eggs that extrude 1 PB (M1). Genes significantly upregulated in M3 eggs appear on the right, those upregulated in M1 appear on the left, and non-significant changes are centered. WT, wildtype.

Next, we compared the DEGs of wildtype eggs to M1 *mos^-/-^*eggs. The M1s look similar to wildtype in that they do not undergo additional divisions. However, we found that the M1s have a very different gene expression pattern with 2,322 DEGs (2,287 upregulated and 35 downregulated; Fig. 7B). Many of the genes were also in cell cycle pathways, including G2/M transition, cell cycle checkpoints, mitotic cell cycle, and APC/C mediated degradation of cell cycle proteins (Fig. 7C). These results suggest that despite their morphological similarities to wildtype eggs, even the *mos^-/-^* eggs with one polar body had increased gene expression, including of cell cycle regulators.

Finally, we asked if the gene expression differs based on whether the eggs remain with one polar body or undergo additional divisions. We compared the M1 eggs to the M3 parthenotes and found very few genes that had increased expression in the M3s (Fig. 7D). There were some genes with decreased expression in the M3s compared to the M1s, including genes involved in the zona pellucida, prevention of polyspermy, interaction with cumulus cells, and regulation of the reproductive process. We found these results surprising because we expected that the M3 eggs would have an increased number of DEGs in comparison to the M1s to be able to enter into the additional divisions. These results demonstrate that even *mos^-/-^* eggs with one polar body have *de novo* transcription, not just the ones that undergo additional divisions. Overall, our results suggest that MOS prevents the eggs from both undergoing parthenogenetic activation and additional cell divisions, while also preventing entrance into a cell cycle state that allows for new transcription.

## DISCUSSION

MOS has long been known to be a regulator of metaphase II arrest. Initial analysis of the *Mos* knockout mice showed that a subset of eggs failed to maintain the metaphase II arrest and instead undergoes parthenogenetic activation and abnormal cell divisions (Colledge *et al*., 1994; Hashimoto *et al*., 1994). Furthermore, approximately one-third of *mos^-/-^*mice develop germ cell tumors over time, suggesting that the activated eggs remain in the ovary, instead of being ovulated, and then develop into tumors (Colledge *et al*., 1994; Hashimoto *et al*., 1994; Hirao and Eppig, 1997). These initial studies led us to further analyze the consequences of the loss of MOS during meiosis and in parthenogenesis. Our results uncovered three major findings. First, we find a novel role for MOS in temporally regulating meiosis I chromosome segregation. Second, we find that new transcription is needed for *mos^-/-^* eggs to undergo additional cell divisions beyond anaphase II, but not for parthenogenetic activation and completion of meiosis II. Third, we show that MOS prevents eggs from exiting metaphase II and entering a cell-cycle state that permits new gene expression, even in the eggs that extrude only one polar body.

To understand the role of MOS in meiotic cell cycle regulation, we used live imaging to compare wildtype and *mos^-/-^* oocytes. We found that some *mos^-/-^*oocytes undergo abnormal chromosome segregation attempts in which the spindle elongates and chromosomes are initially pulled apart, coinciding with membrane morphology changes (Figure 1). However, the spindle and chromosomes come back together, and the morphology returns to that of a normal oocyte. This segregation attempt occurs several hours earlier than normal meiosis I timings in a wildtype oocyte and occurs prior to the movement of the spindle to the cell cortex. Previous work has highlighted a role for MOS/MAPK in spindle positioning (Verlhac *et al*., 1996, 2000; Chaigne *et al*., 2013, 2015). One role is through softening the cell cortex in meiosis I by decreasing cortical tension and through cortical F-actin thickening (Chaigne *et al*., 2013, 2015). In the attempted chromosome segregation events that we observed, the spindle is centrally positioned, suggesting that the coordination between chromosome segregation and the cytoskeletal changes for polar body extrusion is disrupted. The second chromosome segregation event that leads to polar body extrusion then occurs approximately 9 hours later. Because this phenotype only occurs in a subset of oocytes, we propose that MOS may function redundantly with other cell cycle regulators to temporally regulate meiotic chromosome segregation events and to coordinate chromosome segregation and cell division.

Our time-lapse imaging of meiosis II provided additional insight into the behavior of *mos^-/-^* eggs as they escaped the metaphase II arrest. Notably, among the eggs that remained with one polar body, a fraction escaped metaphase II between 0-4hrs and entered an interphase-like state, without extruding a polar body (Figure 3). The eggs that had two polar bodies or underwent additional divisions, also exited metaphase II within 4hrs and either remained in an interphase-like state or underwent additional cell divisions. These results suggest that only a small fraction of *mos^-/-^* eggs maintained the metaphase II arrest. A previous study showed that *mos^-/-^*eggs can reaccumulate cyclin B1 and securin levels after anaphase I, demonstrating that MOS initially inhibits APC/C (Pasternak *et al*., 2016). However, the levels of cyclin B1 are lower in *mos^-/-^*eggs and they are unable to maintain the arrest (Colledge *et al*., 1994; Hashimoto *et al*., 1994; Madgwick and Jones, 2007; Pasternak *et al*., 2016; Cao *et al*., 2020).

Because cyclin B1 binds and activates CDK1, it will be interesting to test how differing levels of cyclin B1 in individual eggs contribute to the variability in the phenotypes.

To test whether new transcription may cause *mos^-/-^* eggs to undergo parthenogenetic activation and additional divisions, we monitored eggs after the addition of a transcriptional inhibitor. Interestingly, we found a substantial decrease in the number of eggs that underwent additional divisions (Figure 2). The percentage of eggs with one polar body significantly increased, while the number with two polar bodies remained similar to *mos^-/-^* eggs without the transcription inhibitor. These results suggest that new transcription is required for the additional cell divisions, but not for parthenogenetic activation.

However, because the number of eggs with two polar bodies did not increase with the inhibition of transcription, new transcription may contribute to parthenogenetic activation in the subset of eggs that would have otherwise undergone additional divisions.

Finally, we used single egg or single parthenote sequencing to identify differentially expressed transcripts depending on both the phenotypic outcome and genotype of the eggs (Figure 4-7). We had expected that only the *mos^-/-^*eggs that underwent additional divisions would have undergone new transcription, while the ones with one polar body or two polar bodies would not. Furthermore, we predicted that the DEGs in the *mos^-/-^* eggs would be similar to that of the chemically activated wildtype eggs because those eggs activate and enter into embryo-like divisions. To our surprise, we found that new transcription occurred in all three categories of *mos^-/-^*eggs, even those with only one polar body (Figure 7). Additionally, *mos^-/-^* eggs and chemically activated wildtype eggs had a very different transcriptional profile (Figure 6). The *mos^-/-^*eggs were found to maintain maternal transcripts, express embryonic genes, and express genes that were normally low in the embryo (Figure 4-5). This discordant expression pattern likely contributes to the additional divisions. These results highlight the important role of MOS in not only maintaining the metaphase II arrest, but also in maintaining the chromatin state that prevents new gene expression.

Overall, our results have implications for understanding infertility. Women with biallelic MOS variants have presented in fertility clinics with eggs that, when fertilized, undergo abnormal early embryonic divisions (Zhang *et al*., 2021, 2022; Jiao *et al*., 2022). The resulting embryos are not viable due to early embryonic arrest and fragmentation. Our results from *mos^-/-^* mice suggest that the presence of the maternal transcripts combined with abnormal new transcription in *mos^-/-^*eggs, even those with one polar body, would result in an egg that is not competent for normal embryo formation after fertilization. Therefore, proper MOS signaling is important to prevent recurrent early pregnancy loss.

Thus, these results provide new insights into the cause of infertility in women with MOS variants and variants of upstream regulators and downstream targets, suggesting new directions to search for therapeutic interventions.

## METHODS

### Mouse strains, husbandry, and genotyping

All animal work in this research was performed at Geisel School of Medicine at Dartmouth College and approved by the Institutional Animal Use and Care Committee at Dartmouth College (Protocol# 00002303). Sexually matured female wildtype and *mos* knockout mice (8-11 weeks old) were used for all experiments. The original heterozygous female and male mice were obtained from the Jackson Laboratories (JAX stock #002723), from the original publication in which a neomycin resistant cassette was introduced in the first exon, creating an amber stop codon upstream of *Mos* kinase domain in a C57BL/6J background (Colledge *et al*., 1994). Experimental mice were obtained by mating heterozygous females to homozygous or heterozygous males. Genotyping was performed using the forward primer 5’-TCAGCTGCAGAGAACAACTGA-3’ in combination with these two reverse primers: 5’-GTGTACGTGCCCCCTATGTG-3’, to detect the presence of the wildtype *Mos* allele, or 5’-GCCAGAGGCCACTTGTGTAG-3’, to identify the neomycin resistance cassette upstream of the *Mos* kinase domain. Mice were housed in a room programmed 12-hour dark/light cycle and constant temperature (72+/-3°F), humidity (30-70%) and with food and water provided *ad libitum*.

### Mouse oocytes isolation and maturation

Prior to each experiment, females were injected intraperitoneally with 5 I.U. of pregnant mare serum gonadotropin (PMSG; BioVendor, RP1782725000) 48h prior to oocyte collection. Ovaries were dissected and prophase I-arrested oocytes were collected using minimal essential medium (MEM) with Earle’s salts (Millipore Sigma, M0268), 0.1g pyruvate (Millipore Sigma, P4562), 1mL gentamycin (Millipore Sigma, G1272), 25mL 1M HEPES pH 7.3 (Millipore Sigma, H3784), 3g polyvinylpyrrolidone pH 7.3 (Millipore Sigma, P2307), and with 2.5μM milrinone (Millipore Sigma, M4659) to prevent meiotic resumption. Oocytes were then cultured in Chatot, Ziomek, and Bavister (CZB) media, which is made up of 81.6mM NaCl (Millipore Sigma, S5886), 4.8mM KCl (Millipore Sigma, P5405), 1.2 mM KH_2_PO_4_ (Millipore Sigma, P5655), 1.2 mM MgSO_4_·7H_2_O (Millipore Sigma, M7774), 0.27 mM Pyruvic acid (Millipore Sigma, P4562), 1.7 mM CaCl_2_·2H_2_O (Millipore Sigma, C7902), 30.8 mM DL-Lactic acid (Millipore Sigma, L7900), 7mM Taurine (Millipore Sigma, T0625), 0.1mM EDTA (Millipore Sigma, E5134), 25mM NaHCO_3_ (Millipore Sigma, S5761), 0.1% gentamicin (Millipore Sigma, G1272), 0.3% BSA (Millipore Sigma, A4503) without milrinone, which allowed for meiotic progression. *In vitro* maturation of oocytes took place in a humidified incubator programmed to 5% CO_2_ and 37°C. Each experiment was independently performed at least twice, with multiple control and test mice included in each repetition.

To chemically activate eggs, wildtype oocytes were matured in CZB for 15-16h after GV release. The eggs were washed and incubated in Ca^2+^-free CZB containing 10mM strontium chloride (Millipore Sigma, Aldrich 25521) for 2h (Jellerette *et al*., 2000). The strontium chloride was then washed in CZB and the eggs were kept in CZB for imaging. Activation was verified by the presence of a pronucleus and/or a 2^nd^ PB.

### Drug additions

To inhibit transcription, actinomycin D (MedChemExpress, HY-17559) was reconstituted in DMSO and added to *mos^-/-^* oocytes 12h after GV release to a final concentration of 1μM actinomycin D in CZB. The actinomycin D stock solution was aliquoted and stored at-20°C for no longer than 1 month, and the final concentration was selected according to previous studies (Zhu *et al*., 2023; Udagawa *et al*., 2024).

### Staining of mouse oocytes/eggs and live imaging

SirDNA and SPY555-tubulin (Cytoskeleton, CY-SC007 and CY-SC203) were reconstituted in DMSO. These dyes were added (100-150nM each) to the oocytes in maturation media (CZB) 45min-1h before imaging and were present throughout imaging. For live imaging, we used a spinning disk CSU-SoRa confocal microscope equipped with a humidified box programmed to 5% CO_2_ and 37°C. Oocytes and eggs were imaged in 250μL of CZB using a chambered cover glass (Cellvis, C18-1.5H) and a 40x Plan Apochromat λS 1.15 numerical aperture water-immersion objective. Groups across all experimental repetitions were imaged on the same microscope using identical imaging settings. Tubulin and DNA were visualized with exposure times of 200ms and 30ms, using neutral-density filters that transmitted 25% and 5% of the light intensity, respectively. During time-lapse imaging, 13 z stacks of 4μm separation were acquired at 20-min intervals for 24h.

### Immunofluorescence and image acquisition

Wildtype and *mos^-/-^* oocytes, eggs, and parthenotes that were immunostained for DNA and tubulin were first fixed using 2% PFA in phosphate-buffered saline (PBS) for 20min at room temperature. They were then kept in blocking solution (0.3% BSA, 0.01% Tween-20, 0.02% NaN_3_, 1xPBS) for no longer than 48h. These eggs were incubated in permeabilization solution (0.3% BSA, 0.1% Triton X-100, 0.02% NaN_3_, 1xPBS) for 20min at room temperature and washed 3 times in blocking solution, each for 10min. To visualize tubulin, eggs were transferred to blocking solution containing alpha-tubulin (11H10) rabbit monoclonal antibody conjugated to AlexaFluor® 488 (Cell Signaling Technology, 5063S; 1:100 dilution) in a humidified chamber for 1h at room temperature. The eggs were washed 3 times in blocking solution, 10min per wash, and mounted on 5μL of Vectashield (Vector Laboratories, H-1000) containing 4,6-Diamidino-2-Phenylindole Dihydrochloride (DAPI; Life Technologies, D1306; 1:170) to detect DNA. These images were captured using a spinning disk CSU-SoRa confocal microscope and a 60x Plan Apochromat λD 1.42 numerical aperture oil-immersion objective. Tubulin and DNA were visualized with exposure times of 700-900ms, using neutral-density filters that transmitted 20-25% of the light intensity each. Images shown are maximum projects of >100 images through 0.3μm z stacks.

Wildtype and *mos^-/-^* oocytes, eggs, and parthenotes that were immunostained for DNA and F-actin were first fixed using 4% PFA in PBS for 30min at room temperature. They were then kept in blocking solution for no longer than 48h. They were incubated in permeabilization solution for 20min at room temperature and washed 3 times in blocking solution, each for 10min. They were incubated in 1% BSA (in PBS) for 1h at room temperature. To visualize F-actin instead, they were transferred to blocking solution containing phalloidin staining (Molecular Probes; 1:20 dilution) in a humidified chamber for 1h at room temperature. These eggs were then washed 3 times with blocking solution, 10min each, and mounted on 5μL of Vectashield (Vector Laboratories, Cat# H-1000) containing DAPI (Life Technologies, Cat# D1306; 1:170) to detect DNA. These images were captured using a spinning disk CSU-SoRa confocal microscope and a 60x Plan Apochromat λD 1.42 numerical aperture oil-immersion objective. F-actin and DNA were visualized with exposure times of 700-900ms, using neutral-density filters that transmitted 20-25% of the light intensity each. Images shown are maximum projections of >100 images through 0.5μm z stacks.

### Single-cell RNA sequencing libary preparation

Single-cell RNA-sequencing (scRNA-seq) libraries were generated using SMART-Seq2 protocol (Trombetta *et al*., 2014). Briefly, cDNA was reverse transcribed from single cells using Superscript III reverse transcriptase (Thermo Fisher Scientific, 18080093) and whole transcriptome amplification (WTA) was performed. WTA products were purified using the Agencourt AMPure XP beads (Beckman Coulter) and used to prepare paired-end libraries with Nextera XT (Illumina). Single cells were pooled and sequenced on a NextSeq2000 sequencer (Illumina) using a P1 100 cycle Kit (v2.5).

### Single-cell RNA sequencing. Analysis and statistics

The high-performance computing (HPC) cluster at Dartmouth was used to perform data pre-processing steps, including data filtering, alignment and generating count matrices. Raw sequencing data were downsampled to 10 million reads per sample to make sequencing depth uniform across samples. The downsampled oocytes were aligned to the GRCm38 genome using STAR aligner (Dobin *et al*., 2013). The final dataset consisted of 108 oocytes with alignment rate above 70%. The count matrices were generated using HTSeq package (Anders *et al*., 2015). These count matrices were further analyzed in R (v.4.1.0) using the Seurat (v5) package (Hao *et al*., 2021). Cells expressing less than 200 genes were excluded from further analysis and genes were filtered to contain only those that were protein coding. To account for the effects of integrating data from multiple sequencing runs, the Harmony package was used to correct for batch effect (Korsunsky *et al*., 2019). Top 10 differentially expressed features were identified based on Type (mutant, chemically activated or WT), or Class (M1, M2, M3, chemically activated or WT). Differentially expressed genes were identified through DESeq2 with an adjusted P value cutoff of 0.05 (Love *et al*., 2014). Heatmaps of top differentially expressed genes were generated using the scaled data. Gene ontology (GO) and pathway analysis were performed using web-based portal Metascape for GO biological process and GO molecular function, using the differentially expressed genes identified in the analysis (Zhou *et al*., 2019). The distribution of aligned reads was examined using Integrative Genomics Viewer (IGV) (Thorvaldsdóttir *et al*., 2013).

## ACKNOWLEDGEMENTS

We thank the Lacefield Lab for reading the manuscript. RNA sequencing was carried out in the Genomics and Molecular Biology Shared Resource at Dartmouth (RRID:SCR021293) which is supported by NCI Cancer Center Support Grant 5P30CA023108 and NIH S10 (1S10OD030242) awards. SL is supported by the bioMT through NIH NIGMS grant P20-GM113132. BAG is supported in part through the Geisel School of Medicine at Dartmouth’s Center for Quantitative Biology through a grant from NIH NIGMS P20GM130454.

---

**Figure S1.**
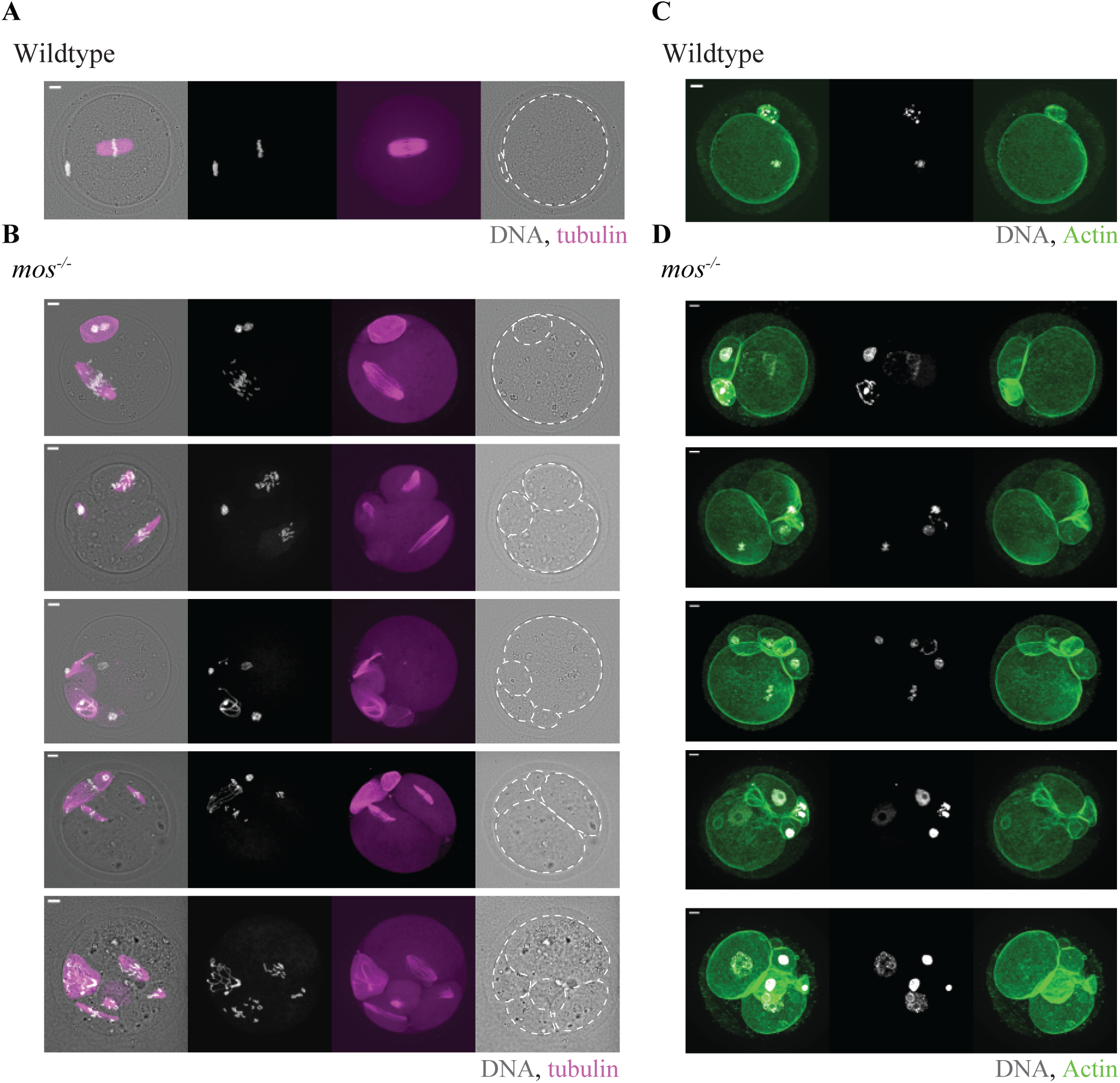
***mos^-/-^*eggs exhibit different numbers of polar bodies and cells**. **(A-D)** Representative immunofluorescence images with the indicated genotype showing localization of DNA (white) with either tubulin (magenta) (A-B) or F-actin (green) (C-D). Eggs were fixed 24h after meiotic resumption. Scale bars, 10µm. WT, wildtype.

**Figure S2.**
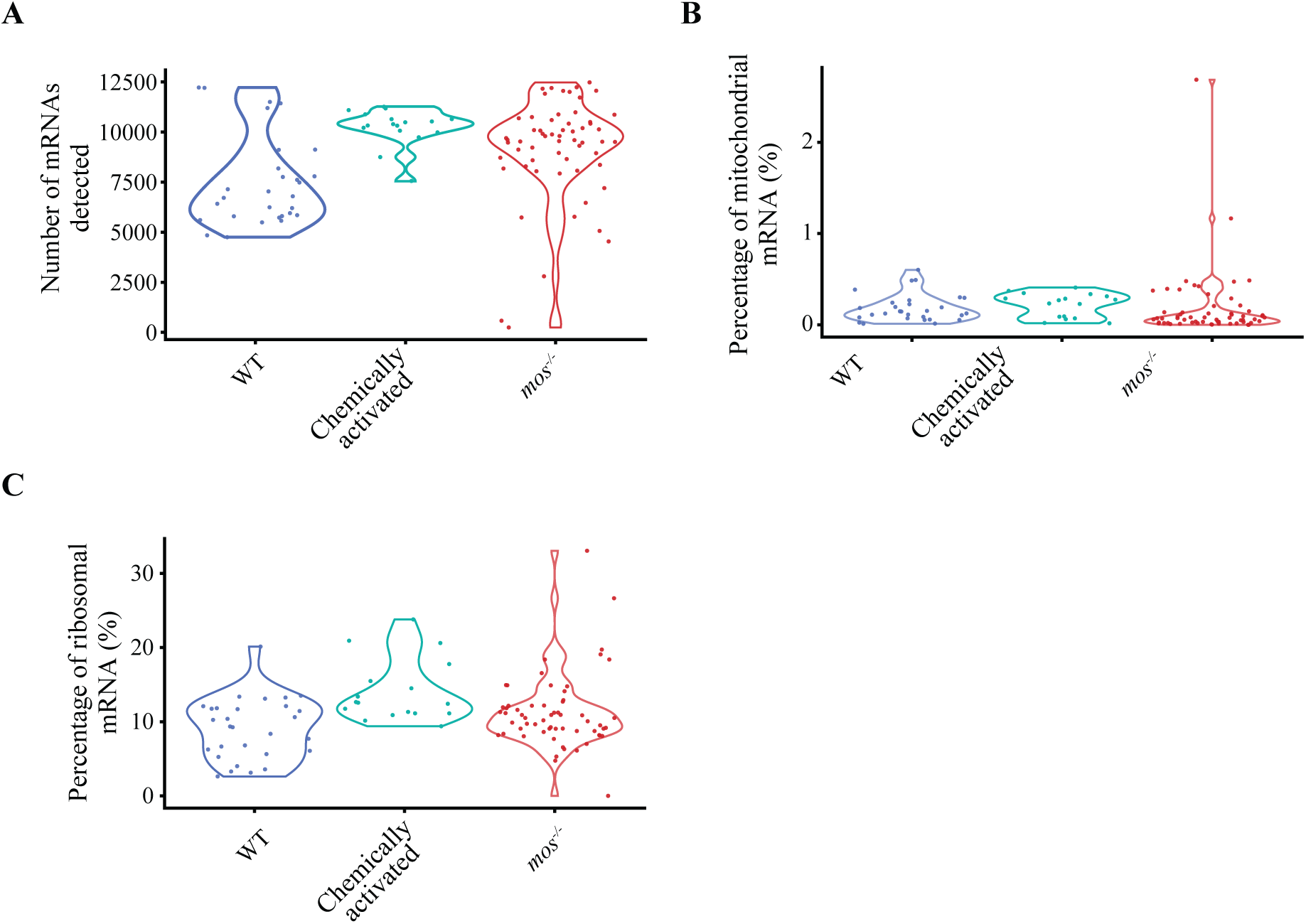
QC metrics for scRNAseq analysis. (A-C) Violin plots representing the number of mRNAs detected per cell (A), percent of mitochondrial mRNA (B), and percent of ribosomal mRNA in wildtype, chemically activated, and *mos^-/-^*eggs used for scRNAseq. WT, wildtype.

## Notes

### Competing Interest Statement

The authors have declared no competing interest.

